# Induced antigen-binding polyreactivity in human serum IgA

**DOI:** 10.1101/2021.09.29.461938

**Authors:** Ekaterina N. Gorshkova, Maxime Lecerf, Irina V. Astrakhantseva, Ekaterina A. Vasilenko, Olga V. Starkina, Natalya A. Ilyukina, Petya A. Dimitrova, Jordan D. Dimitrov, Tchavdar L. Vassilev

## Abstract

Previous studies have shown that polyreactive antibodies play an important role in the frontline defense against the dissemination of pathogens in the pre-immune host. Interestingly, antigen-binding polyreactivity can not only be inherent, but also acquired post-translationally. The ability of individual monoclonal IgG and IgE antibodies to acquire polyreactivity following contact with various agents that destabilize protein structure (urea, low pH) or have pro-oxidative potential (heme, ferrous ions) has been studied in detail. However, to the best of our knowledge this property of human IgA has previously been described only cursorily. In the present study pooled human serum IgA and two human monoclonal IgA antibodies were exposed to buffers with acidic pH, to free heme or to ferrous ions, and the antigen-binding behavior of the native and modified IgA to viral and bacterial antigens were compared using immunoblot and ELISA. We observed a dose-dependent increase in reactivity to several bacterial extracts and to pure viral antigens. This newly described property of IgA may have therapeutic potential as has already been shown for pooled IgG with induced polyreactivity.

## Introduction

The enormous diversity of antibody specificities is the result of complex genetic recombination and mutational processes. Epigenetic mechanisms can increase this diversity still further [1]. Naturally polyreactive antibodies, referred to also as multispecific, promiscuous or degenerate, are present in the blood of all healthy individuals [2]. While most IgM antibodies are known to be polyreactive, only a portion of IgG and IgE immunoglobulins possess this property. Scientific proof for the existence of polyreactive antibodies first came when it was demonstrated that some monoclonal antibodies could bind more than one structurally different antigen. The molecular mechanisms of polyreactive antigen binding are discussed in detail elsewhere [3].

Natural polyreactive antibodies play an important role as a frontline defense against invading pathogens, in the maintenance of immune homeostasis, in the clearance of senescent and damaged cells, and etc. [4]. Twenty years ago it was shown that in addition to innate (natural) polyreactivity, some immunoglobulins can acquire this property after exposure to chaotropic agents, buffers with low pH, or high salt solutions [5]. Later it was demonstrated that the same effect can be achieved after exposure of human pooled IgG to pro-oxidative agents such as heme and ferrous ions [6,7]. We hypothesized that this induced polyreactivity may be observed not only *in vitro*, but also *in vivo* – at sites of inflammation where various aggressive protein-modifying molecules are released. Indeed, it was later demonstrated that induction of polyreactivity through exposure of IgG molecules to protein-modifying agents is not only in vitro phenomenon, but can also occur *in vivo* [8]. Free heme and ferrous ions are known to be released *in vivo* at sites of inflammation, trauma, as result of intravascular hemolysis, and etc. The ability of heme to induce IgG antigen-binding polyreactivity has been reported by J. A. McIntyre *et al*. [6]. Moreover, exposure of immunoglobulins to histones can also trigger their polyreactivity [9].

Our previous studies have shown that the exposure of human IgG to heme, to ferrous ions or to buffers with acidic pH results in the induction of different degrees of polyreactivity [1,7,10,11]. Pooled human IgG for intravenous administration (intravenous immunoglobulin, IVIg) with an enhanced ability to bind additional antigens has therapeutic potential. While the infusion of native (untreated) pooled human IVIg has no beneficial effect in mice with experimental sepsis, the same dose of ferrous ions-treated IVIg increases the animals’ survival significantly [7]. It was also shown that IVIG can suppress the induction of diabetes, possibly through the induction of T-regulatory cells, and it was hypothesized that the effect of IVIG after binding to heme was enhanced [12]. Recently we have also shown that a monoclonal IgE antibody exposed to heme can acquire additional antigen-binding polyreactivity [13].

More serum and secretory IgA are produced per day in the human body than all antibodies of the other immunoglobulin classes taken together. Much is known about natural polyreactive secretory IgA (sIgA). Immune protection against microorganisms in the digestive tract is primarily provided by sIgA. Moreover, this occurs both in adults due to their own sIgA, and in infants due to sIgA obtained from colostrum [14–17]. Also, our group confirmed the possibility of acquiring the properties of polyreactivity by secretory immunoglobulins A, and these changes were multidirectional, depending on the inducing agent [18]. While the role of secretory IgA in health and disease is quite well understood, that of the serum IgA is not so well elucidated. Recent studies have shown that serum IgA plays a role in the immune response, which differs from the role of secretory IgA in intestinal immunity. As some studies have described, the monomeric form and the lack of a secretory component allows serum IgA to bind Fcα receptor I (FcαRI) of myeloid cells, thereby causing reactions such as the release of cytokines, phagocytosis and the formation of extracellular neutrophil traps (NET) [19,20]. It was also shown that the effector functions utilized by IgA depends on the subclass to which it belongs [21]. Much is still not clear about serum IgA, including whether it is possible to induce its cryptic polyreactivity following exposure to different in vivo relevant triggers.

The aim of our study was to formally prove the ability of heme, low pH buffers and Fe(II) ions to induce polyreactivity in human serum IgA.

## 2. Materials and Methods

### 2.1. Serum IgA preparations

As a source of polyclonal human serum immunoglobulin A (IgA) we used a complex immunoglobulin preparation (RPC Microgen, Russia), which is a lyophilized mixture of normal human immunoglobulins [IgG + IgM + IgA] isolated from human plasma of donors tested for the absence of antibodies to the human immunodeficiency virus (HIV -1, HIV-2), hepatitis C virus and hepatitis B virus surface antigen. Isolation of IgA from a mixture of immunoglobulins was carried out using the agarose-immobilized Jacalin (Thermo Fisher Scientific, USA). For this purpose, 4 mL of 50% gel slurry were loaded on a gravity column. After sedimentation the packed bed was equilibrated with 10 mL of PBS. The lyophilized immunoglobulins preparation was diluted to a concentration of 15 mg / mL, loaded onto the columns in a volume of 3 mL and the unbound fraction was washed with 10 mL PBS. IgA antibodies were eluted with 10 mL elution buffer (0.1M alpha-D-galactose in PBS) in 2 mL fractions. IgA-containing fractions were pooled and dialyzed against PBS. Concentrations were estimated by spectrophotometry. Serum IgA, exposed to low pH during its fractionation was obtained from the biotechnology company Imtek (Moscow, Russia). Its exposure to heme was performed as described previously [22]. The exposure to pro-oxidative ferrous ions was carried out as follows: serum IgA (5 mg/mL) was incubated (at 4 °C for 30 min) with freshly prepared 1 mM solution of FeSO_4_ in PBS-T buffer, containing 137 mM NaCl, 2.7 mM KCl, 10 mM Na_2_HPO4, 1.8 mM KH_2_PO4, and 0,1% Tween 20 (all from Sigma-Aldrich, USA). Next, the IgA solution was dialyzed for 120 min. at 4°C against 1x phosphate– buffered saline (PBS) containing 5 μM ethylenediaminetetraacetic acid (EDTA) and finally against 1x PBS without EDTA (4°C, 24 h). The following buffers were used to evaluate the ability of low pH conditions to induce IgA polyreactivity: TBS (pH 7.0) and two glycine buffers (with pH 2.6 and with pH 4.0).

### 2.2. Cloning of monoclonal antibodies

Variable regions of the heavy and light chains of mAb21 and mAb72 originated from a previously described repertoire of human antibodies obtained by single-cell PCR from synovial tissue of rheumatoid arthritis patients [23]. Variable regions were cloned in IgA1 expression vectors by the In-Fusion technology (Takara) as per manufacturer’s protocol. The expression vectors were kindly provided by Dr. Hugo Mouquet (Institute Pasteur, Paris France) [24].

### 2.3. Expression and purification of monoclonal antibodies

For each antibody, expression vectors for the heavy and light chain were co-transfected in the ExpiCHO cell line (Thermo Fisher Scientific, USA) using the manufacturer’s protocol. In brief, 2.5 mL of a 6×10^6^ cells/mL suspension were co-transfected with a total of 2.5 μg plasmid DNA (1:1 molar ratio of heavy and light chain expression vectors). Transfected cells were incubated for 8 days at 37 °C and 8% CO_2_ with shaking (225 rpm). The supernatants were harvested by centrifugation at 3000 x g for 30 minutes at 4 °C, filtered through a 0.22 μm syringe filter and diluted with 1 volume of PBS. For each antibody, 4 mL of 50 % gel slurry of agarose-immobilized Jacalin (Thermo Fisher Scientific, USA) were loaded on a gravity column. After sedimentation the packed bed was first equilibrated with 10 mL of PBS. Diluted supernatants were loaded onto the columns and the unbound fraction was washed with 10 mL PBS. IgA antibodies were eluted with 10 mL elution buffer (0.1M alpha-D-galactose in PBS) in 2 mL fractions. IgA-containing fractions were pooled and dialyzed against PBS. Concentrations were estimated by spectrophotometry.

### 2.4. Preparation of antigen extracts for Western blotting

*E. coli* (BL21) and *S. aureus* (2879 M) were cultured on an orbital shaker in sterile LB liquid medium containing 10 g Tryptone, 10 g NaCl, 5 g yeast extract (all from Sigma-Aldrich, USA) prepared in distilled water up to 1 L, (pH was adjusted to 7.5) for 16–18 hours at 37 °C, then centrifugated (at 14 000 rpm for 5 min.). After removal of the supernatant the pellet was resuspended in 5 mL of cell lysis buffer containing 30 mM NaCl, 50 mM, TRIS (pH 6.8), 5 % glycerol, and 0.5 % Triton X-100 (all from Sigma-Aldrich, USA), homogenized by sonication and centrifugated for 15 min at 15000 rpm. The lysate supernatants were used for Western blot analysis.

Cells from the human cancer cell line Colo205 were cultivated in complete RPMI-1640 medium (Thermo Fisher Scientific, USA) supplemented with 2 mM L-glutamine, antibiotics (100 U/ml penicillin, 100 μg/ml streptomycin) and 10 % fetal bovine serum (Thermo Fisher Scientific, USA). The cells were detached from the culture flask using 0.25 % (w/v) Trypsin/0.53 mM EDTA solution and resuspended (at 4×10^6^ cells/ml) in 0.5 mL of lysis buffer [50 mM TRIS base with pH 8.0, 150 mM NaCl, 1% Triton X-100, 0.5% sodium deoxycholate, and 0.1% SDS, all from Sigma-Aldrich, USA] with constant agitation at 4 °C for 30 min. The cells were then centrifugated (4 °C, 20 min, 21 000 rpm) and the supernatant was aspirated, placed in a fresh tube, kept on ice and used for the Western blot analysis.

### 2.5. Western blotting

The proteins from the lysates were transferred to nitrocellulose membranes (Bio-Rad, USA) using standard Sodium Dodecyl Sulfate-Polyacrylamide Gel Electrophoresis (SDS-PAGE) in TRIS/Glycine buffer [25 mM TRIS base (pH 7.8-8.5), 190 mM Glycine, 0.1 % SDS, and 20 % Ethanol, all from Sigma-Aldrich, USA] for 1.5 h at 400 mA. The nitrocellulose membrane (0.2 μm, Bio-Rad, USA) was blocked with 5% non-fat dried milk in TBST buffer [50 mM TRIS base (pH 7.5), 150 mM NaCl, 0.05 % Tween-20, all from Sigma-Aldrich, USA] for 1 h at room temperature to prevent non-specific binding. The membrane was washed 5 times with TBST (for 5 min.), cut into stripes and incubated with the native or the modified (with heme, Fe(II) or acidic pH) antibodies for 2 h at room temperature and washed with TBST five times. Next, the stripes were incubated with a Goat Anti-Human IgA alpha chain (HRP-conjugated) antibody (ab6858, Abcam, UK) (1:10000), washed with TBST 5 times and twice with TBS, and then incubated in 100 mM TRIS-solution (pH 8.0) containing 0,8 mM iodophenylboronic acid, 1.25 mM luminol and 30% hydrogen peroxide. The stripes were then placed in a digital scanner for chemiluminescent blots (LI-COR Biosciences, UK). The results were analyzed using the LI-COR Biosciences Software Image Studio (LI-COR Biosciences, UK).

### 2.6. Enzyme-linked Immunosorbent Assay (ELISA)

To assess the binding activity of serum IgA to viral antigens 96-well polystyrene plates (Corning® 96-well Clear Polystyrene High Bind Stripwell™ Microplate, Corning, USA) were coated overnight at room temperature with the following antigens: Hepatitis C virus NS3 antigen (HCV), Hepatitis D virus antigen (HDV), and the p24 antigen of human immunodeficiency virus type 2 (HIV) (the antigens were kindly provided by RPC Diagnostic Systems, Russia). After blocking with PBS containing 5 % BSA, the wells were washed 4 times with PBS containing 0.0 5% Tween 20 and incubated with increasing concentrations of native or modified IgA for 1 h at 37°C (in PBS containing 0.05% Tween 20 and 1% BSA). After washing, the plates were incubated with a Goat anti-human IgA alpha chain (HRP-conjugated) antibody (ab6858, Abcam, UK) for 1 h, followed by treatment with 3,3′,5,5′-tetramethylbenzidine (Sigma-Aldrich, USA). Optical density values were read at 450/620 nm. The results were analyzed as the ratio between the optical density of native IgA and IgA exposed to protein-modifying molecules. A ratio equal or higher than 2 was considered as a significant increase in polyreactivity.

## 3. Results

### 3.1. Exposure of serum immunoglobulin A and monoclonal IgA to free heme increases binding to antigens

A variety of pathological processes cause degradation of hemoglobin and myoglobin, and release free heme into the circulation [25]. Several plasma proteins - hemopexin, albumin, and some lipoproteins are able to bind heme and eliminate it from the plasma [26]. However, when the quantity of released heme is above the binding capacities of its scavengers, the residual free heme can bind to immunoglobulin molecules and change the nature of the noncovalent forces responsible for their antigen binding [27]. We observed that the exposure of serum IgA to 0.6 μM and 1.2 μM heme slightly increases its binding to antigens from *E. coli* and Colo205 cells (Figure 1A). Similarly, following exposure to heme the monoclonal antibodies mAb21 and mAb72 recognize novel antigens from *E. coli, S. aureus* and Colo 205 cells. (Figure 1 C, D). Exposure to heme failed to change the ability of serum IgA to recognize and bind to antigens from *S. aureus* (Figure 1A) and did not affect its reactivity to the purified viral antigens tested (Figure 1B). In addition, we wanted to further investigate whether or not heme can induce binding of mAb21 and mAb72 to human self-antigens using ELISA (Supplementary method). Indeed, we found the appearance of new reactivities toward various autoantigens (Supplementary Figure S1).

**Figure 1.**
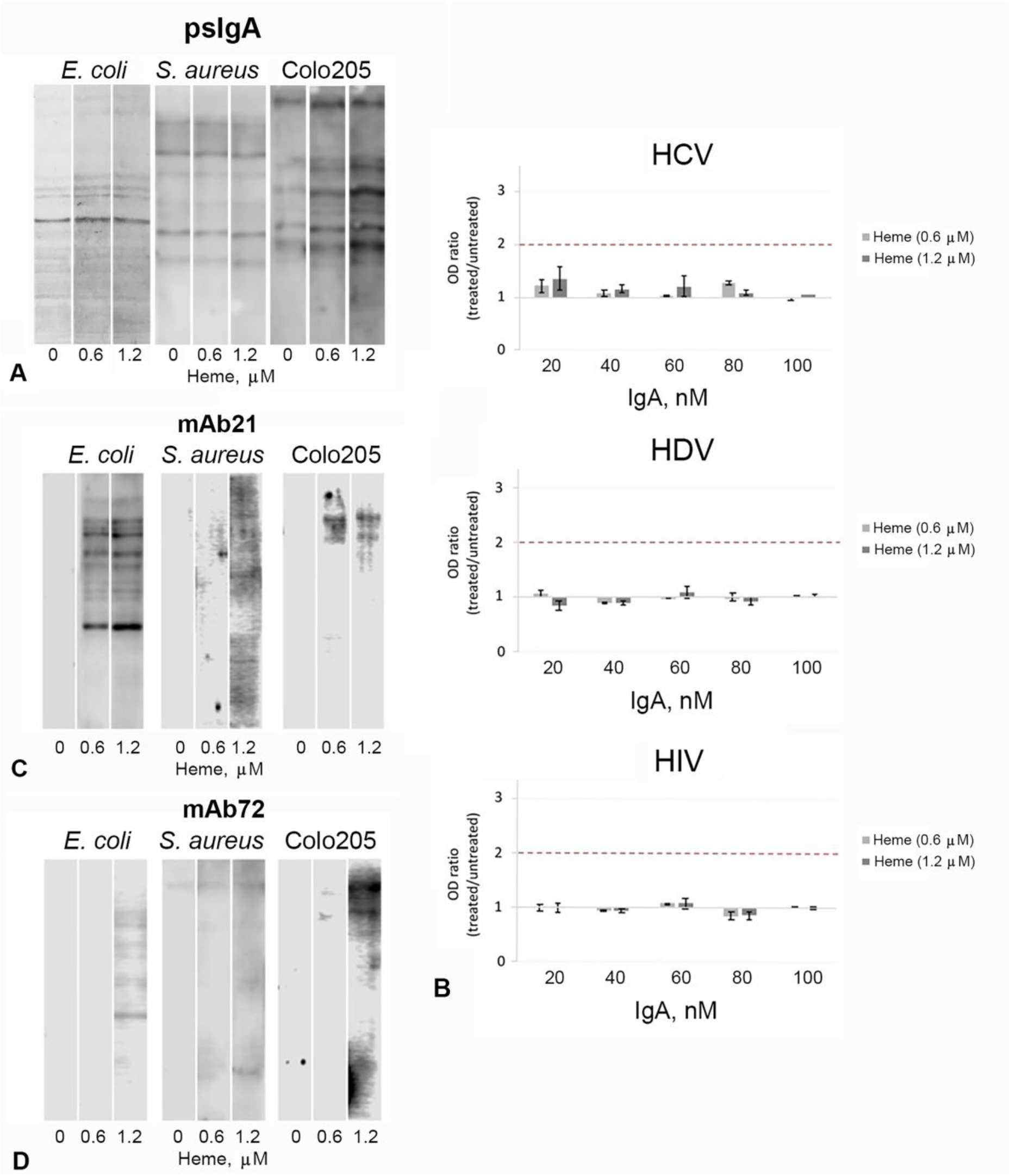
Increased immunoreactivity of human serum IgA and monoclonal antibodies after exposure to heme. (A) Western blot analysis shows that exposure of human serum IgA to heme at concentrations 0.6 μM and 1.2 μM slightly increases its binding to antigens from Colo205 and *E. coli*, which can be judged by an increase in the number and brightness of the bands detected on the membrane. Heme at concentrations 0.6 μM and 1.2 μM do not influence the binding activity of human serum IgA to *S. aureus* antigens: the number of bands did not change after treatment. (B) Heme-treated IgA does not alter the recognition and binding to selected HCV, HDV and HIV antigens as shown by ELISA: the absorbance values for the heme-treated IgA samples were comparable to the values for the untreated IgA throughout the entire concentration range (20 – 100 nM). The data represents mean of four repetitions ±SEM. Exposure of two monoclonal antibodies (C) mAb21 and (D) mAb72 to heme at concentrations 0.6 μM and 1.2 μM allowed their binding to *E. coli, S. aureus* and Colo205 antigens: the absence of bands on the membrane related to non-treated antibodies (0 μM) indicates that the untreated antibodies did not have specificity for these antigens.

### 3.2. Pooled human serum IgA become polyreactiv after exposure to Fe(II) ions monoclonal IgA free heme

It is well known that iron ions possess a potent pro-oxidative potential and it has been shown earlier that the pro-oxidative potential of Fe(II) ions causes enhanced antigen-binding reactivity of IgG antibodies isolated from healthy humans [7]. It has been observed that the exposure of IgG to Fe(II) ions results in a strong increase in their antigen-binding polyreactivity and these modified immunoglobulin preparations have pronounced anti-inflammatory potential in vivo [7,28]. We were interested to explore whether or not Fe(II) ions have the same effect on serum IgA. Indeed, our results show that exposure to Fe(II) ions (concentrations ranging from 25 to 400 μM) increased the binding activity of IgA to *E. coli* and Colo205 antigens in a dose-dependent manner (Figure 2A, B, C). Similar to the results obtained for the heme-treated IgA, we failed to detect a change in the binding activity to *S. aureus* antigens (Figure 2B). Exposure to two different doses of Fe(II) ions (25 and 100 μM) induced binding to viral antigens and reached above the 2-fold threshold (see Materials and Methods) at the highest IgA concentration (100 nM; Figure 2B). Results obtained for monoclonal antibodies shown that incubation with Fe(II) did not change their antigen-binding properties (Supplementary Figure S2).

**Figure 2.**
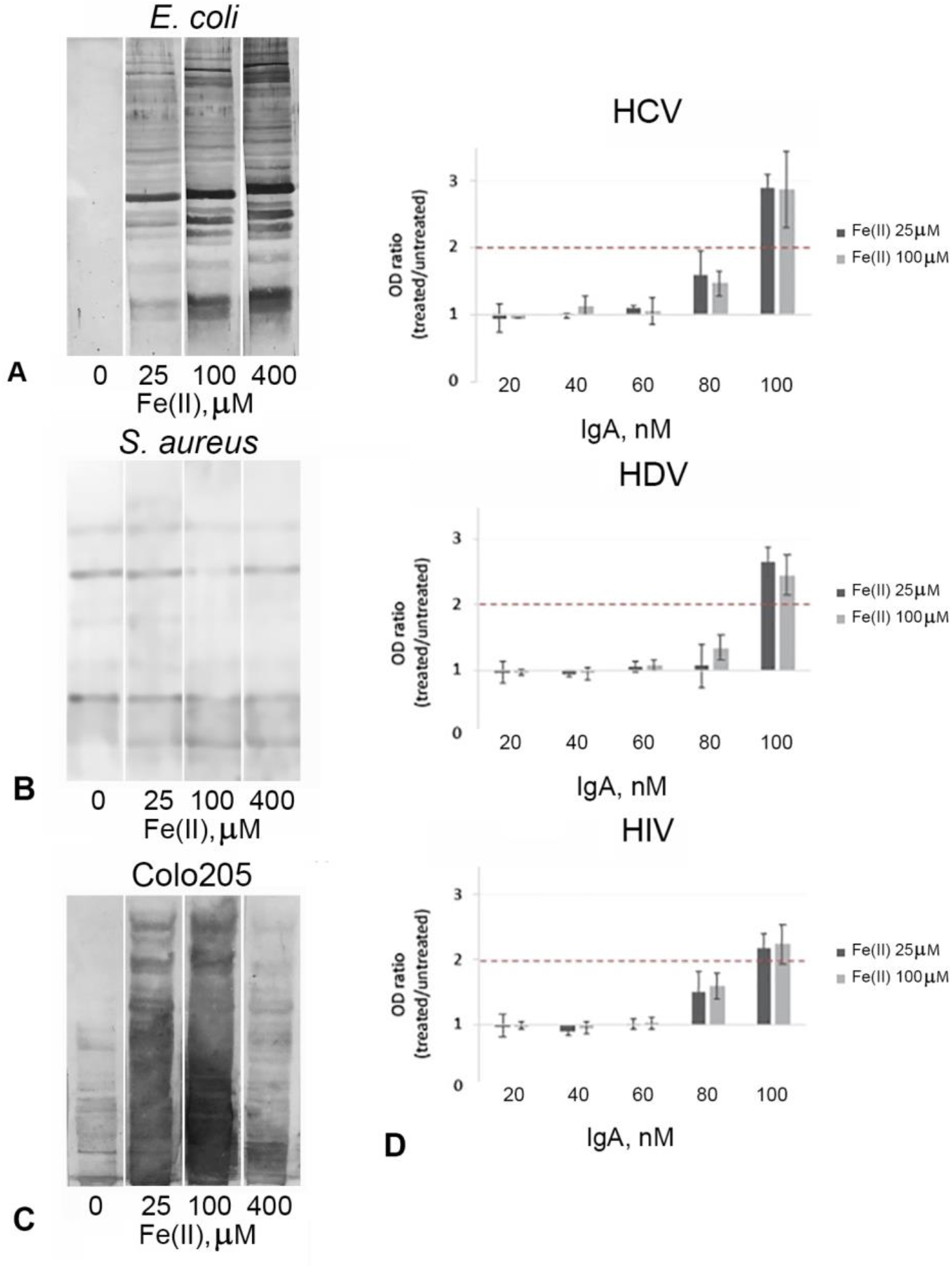
Increased immunoreactivity of human serum IgA after exposure to Fe(II) ions. Western blot analysis shows that the polyreactivity of serum IgA exposed to Fe(II) at concentrations 25 μM, 100 μM and 400 μM induces the binding to *E. coli* (A) and Colo205 (C) antigens, which can be judged by an increase in the number and brightness of the bands detected on the membrane. (B) Fe(II) ions at concentrations 25 μM, 100 μM and 400 μM do not influence the binding activity of human serum IgA to *S. aureus* antigens: the amount of bands did not change after the treatment. (D) Data from the ELISA analyses shows that after treatment with Fe(II) at concentrations 25 μM, 100 μM and 400 μM the same polyclonal IgA acquires enhanced binding ability to all investigated single viral antigens: the absorbance values for the Fe(II)-ions treated IgA samples at concentrations 80 nM and 100 nM were significantly higher than the values for the untreated IgA for HCV, HDV and HIV proteins. The data represents mean of four repetitions ±SEM.

### 3.3. Exposure to buffers with low pH induces IgA polyreactivity

Transient exposure to low pH environment (pH < 4) changes the structure of the IgG molecules and causes an enhancement of IgG antigen-binding polyreactivity [11]. In order to delineate whether or not acidic buffers have a similar effect on the antigen-binding behavior on serum IgA its solutions of different concentrations were exposed to buffers with different pH and then measured the binding activity to bacterial or viral antigens by Western blot or ELISA, respectively. We observed improved binding of IgA after treatment with acidic buffers. Both the repertoire of detected antigens (shown as the number of bands) and the intensity of staining were increased exposure of IgA to low pH (Figure 3A, C). We also observed a modest increase in the IgA antigen-binding ability to individual viral antigens after exposure to an acidic buffer with pH 4.0. Exposure to the buffer with a lower pH (pH=2.6) reduced the antigen binding capacity (Figure 3D). And we did not observe any changes in antigen-binding properties of low pH-treated monoclonal antibodies mAb21 and mAb72 (Supplementary Figure S3).

**Figure 3.**
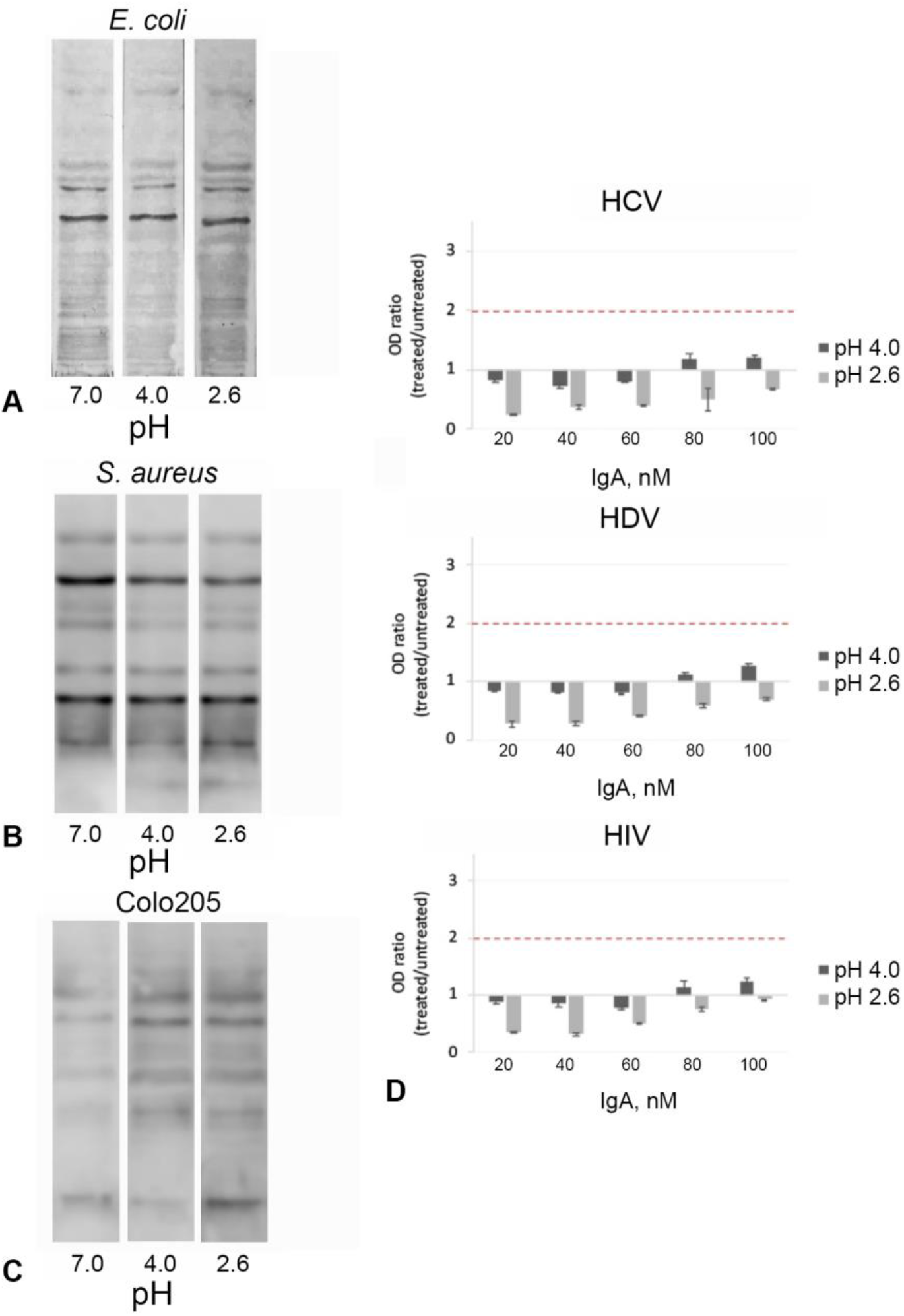
Increased immunoreactivity of human serum IgA after exposure to buffers with low pH. Exposure of human serum IgA to buffers with pH 2.6 but not pH 4.0 results in an increase in the binding to *E. coli* (A) and *S. aureus* (B) and Colo205 (C) antigens, which can be judged by an increase in the number and brightness of the bands detected on the membrane. (D) ELISA results show that after treatment with a buffer with pH 4.0 polyclonal serum IgA at concentrations of 80 nM and 100 nM demonstrates insignificant increase in binding to individual viral antigens. Exposure to the buffer with pH 2.6 reduced its antigen-binding ability throughout the entire concentration range (20 – 100 nM). The data represents mean of four repetitions±SEM.

## 4. Discussion

We have previously described in detail the mechanisms and biological consequences of inducing additional polyreactivity in a human monoclonal IgE antibody (by exposure to heme only [13]), as well as in polyclonal human IgG preparations including one used in the clinic for the treatment of patients (IVIg). The latter has been exposed to free heme [1,27,29], to low pH buffers [5,11] as well as to ferrous ions [28]. The ability to acquire polyreactivity properties has also been shown for secretory IgA [18] The phenomenon of induced polyreactivity has been studied mostly *in vitro*. Furthermore, local concentrations of free heme, pro-oxidative ferrous ions and reactive oxygen species at sites of inflammation and severe trauma could reach levels that are sufficient to modify the antigen-binding behavior of circulating immunoglobulins [8]. IVIg preparations were assayed for the treatment of sepsis and related systemic inflammatory response syndromes [30,31]. However, the higher-quality randomized clinical trials failed to prove any beneficial effect [32]. We have also demonstrated that while the administration of unmodified IVIg had no effect on survival of mice with sepsis and two types of severe aseptic response syndromes, the survival of animals treated by IVIg with additional induced polyreactivity was significantly improved [28]. The present study supports our hypothesis that the exposure of human serum IgA to heme, ferrous ions or to low pH may modulate its antigen-binding specificity in the same manner as reported earlier for IgG. As previously shown for IgG, we could observe that the profile of antigens recognized by serum IgA differ depending on the protein-modifying conditions and lead to certain consequences (“criptic polyspecificity”) [33]. IgA with induced polyreactivity are able to bind to a wide range of antigens, but this effect was not similar for all studied antigens, with the exception of Fe(II) ions treatment, although it failed to induce the polyreactivity of monoclonal antibodies for some antigens (Fig. S2). The effects of heme on monoclonal antibodies were expressed in their ability to bind individual autoantigens. This can probably be explained by the properties of individual monoclonal antibodies. Thus, IgA, like IgG, do not become indiscriminately binding, but have certain preferences. Based on this, it should be assumed that the nature of these changes is similar to that for IgG and might occur by the same mechanisms. As shown earlier, short-term exposure to acidic pH and pro-oxidative Fe(II) ions leads to structural changes in the paratopes of the susceptible immunoglobulin molecules. This chenges are characterized by an increase of the structural dynamics. The enhanced flexibility and better adaptability of paratope might explain the capacity of the modified antibodies to interact with a wider range of epitopes [5,7,11]. The induction of antibodies polyreactivity after exposure to heme has a different molecular mechanism: its interaction with proteins rigidifies their polypeptide chains as demonstrated by kinetic and thermodynamic analyses (described here [34]). But antibodies use heme as interfacial cofactor for binding to new antigens.

Our results suggests that serum IgA with induced polyreactivity may have a therapeutic potential as has been previously reported for IgG [29]. Plasma IgA has potent antimicrobial properties through mechanisms that include neutralizing toxins, activating the alternative complement pathway and phagocytosis and NETs by neutrophils, and some IgA preparations have already been clinically tested [35]. We believe that additional modification with an increased ability to bind new antigens may increase the therapeutic potential of such drugs. Also, the results obtained in this study show that the peculiarities of obtaining and purifying immunoglobulin A should also be taken into account. The method of choice for the rapid purification of immunoglobulins of human and animal origin is based on the specific binding of its molecules to immobilized Protein A and Protein G. The elution step is performed by washing of the respective immunoaffinity columns with low pH buffer. Most of the commercially available purified human serum IgA and secretory IgA are produced in this way. Our studies have shown that this treatment causes enhancement of IgA antigen-binding polyreactivity. The subsequent use of these modified immunoglobulin molecules in different experiments will inevitably result in biased results (Supplementary Figure S4). This suggests that this approach should not be used and should perhaps be replaced by milder techniques (e.g. using the gentle Ag/Ab elution buffers).

In conclusion, it should be noted that it is important to understand that the role of serum IgA in the immune response has not been fully appreciated and the results obtained indicate that the ability of serum IgA to change antigen-binding behavior must also be considered. It is known that monomeric IgA plays a significant role in maintaining homeostasis through the FcαRI receptors, inducing immune cells and pro-inflammatory responses resulting in elimination of pathogens and cancer cells [36,37]. At the same time excessive formation of IgA immune complexes can result in heightened neutrophil activation and autoimmune damage [36,38]. Nevertheless, it is important to keep in mind that while expansion of the repertoire of heme-treated antibodies may provide additional protection against pathogens, it may also contribute to the pathogenesis of inflammation and autoimmunity [27].

## Acknowledgments

We are grateful to Tatiana Tchurina, Mariya Balykova and Elizaveta Razzorenova for their contribution to experimental work, Vladislav Mokhonov and Dmitry Novikov for support and assistance in experiment design.

## Author contributions

Ekaterina N. Gorshkova: Formal Analysis, Funding Acquisition, Supervision Project Administration, Visualization, Writing - Original Draft

Maxime Lecerf: Formal Analysis, Data Curation, Investigation, Methodology, Visualization

Irina V. Astrakhantseva: Methodology, Validation, Writing - Review & Editing

Ekaterina A. Vasilenko: Data Curation, Investigation

Olga V. Starkina: Investigation, Resources

Natalya A. Ilyukina: Investigation, Visualization

Milena Leseva: Writing - Review & Editing

Petya A. Dimitrova: Funding Acquisition, Validation, Writing - Review & Editing

Jordan D. Dimitrov: Conceptualization, Methodology, Writing - Review & Editing

Tchavdar L. Vassilev: Conceptualization, Funding Acquisition, Project Administration, Writing - Review & Editing

