## Supplementary materials for "Induced antigen-binding polyreactivity in human serum IgA"

**Supplementary method.** To assess the binding activity of monoclonal antibodies 96-well polystyrene plates NUNC MaxiSorp (Thermo Fisher Scientific) were coated for one hour at room temperature with human proteins [factor IX (LFB laboratories France), histone 3 (Sigma-Aldrich), hemoglobin, myoglobin, insulin, thyroglobulin and cytochrome C (all from TSigma-Aldrich)]. All proteins were diluted to a concentration of 10 µg/mL in PBS. After blocking for 60 min with a 0.25 % solution of Tween 20 in PBS, the plates were incubated with IgA antibodies. Each antibody was first diluted in PBS to a concentration of300 µg/mL, and then either exposed, or not, for 10 min to 2 µM hemin (stock solution in 0.05 N sodium hydroxide) and then diluted to 60 µg/mL in PBS-T. The antibodies were further serially diluted (3-fold) in the plates, from 60 to 0.083 µg/mL and incubated with coated proteins for 120 min at RT. For assessment of the binding of each recombinant antibody in its native and modified forms to the studied proteins, the plates were first washed5 times with PBS-T and then incubated for 60 min at RT with an HRP-conjugated anti- human IgA (heavy chain-specific, polyclonal, Southern Biotech) diluted 3000× in PBST. Following washing with PBS-T, the immunoreactivity of the antibodies was revealed by theaddition of 50 µL of the peroxidase substrate, o-phenylenediamine dihydrochloride (Sigma-Aldrich). Optical density was measured with a microplate reader (Infinite 200 PRO, Tecan)at 492 nm after addition of 25 µL 2M HCl.

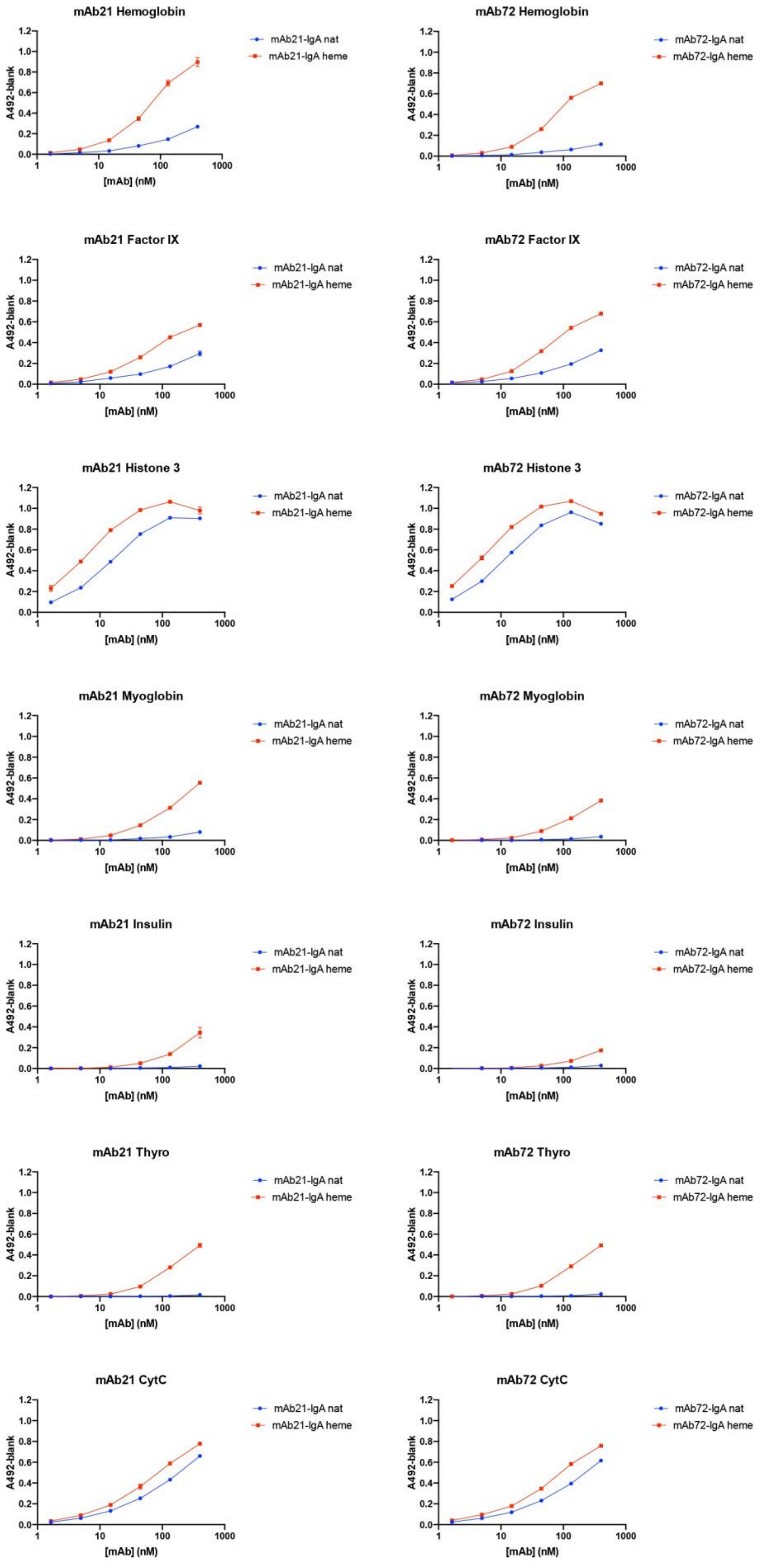


**Figure S1.** Binding activity of monoclonal antibodies mAb21 and mAb72 to human proteins: factor IX, histone 3, hemoglobin, myoglobin, insulin, thyroglobulin (Thyro) and cytochrome C (CytC)

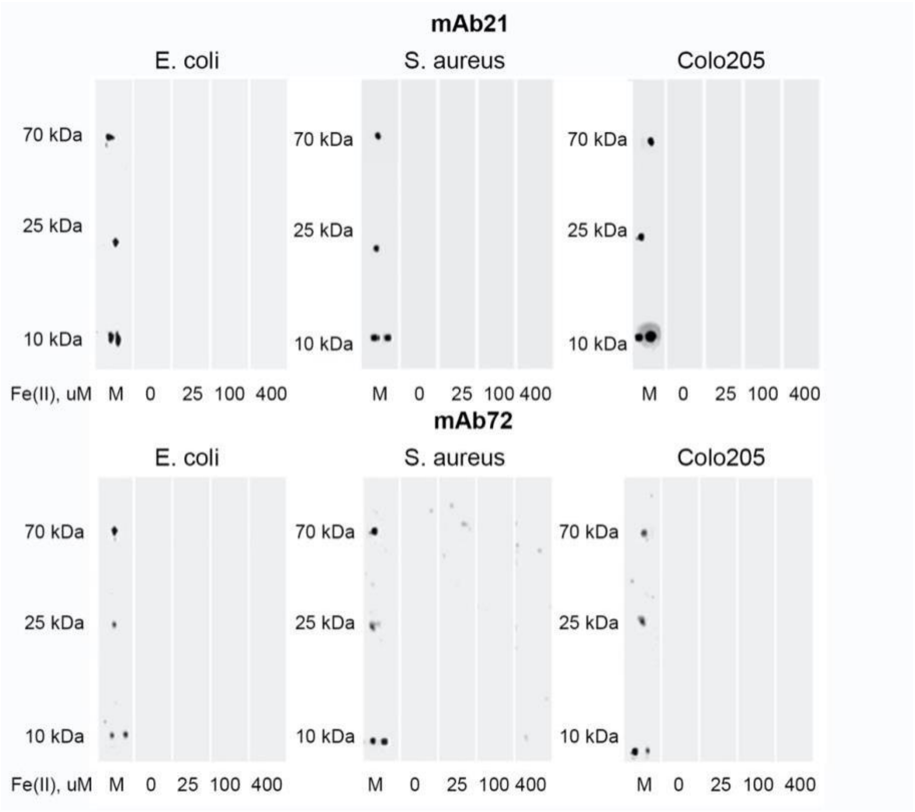


**Figure S2.** Western blot analysis of monoclonal IgA (mAb21 and mAb27) with extracts of *E. coli*, S. *aureus* и *Colo 205.* Antibodies was treated with Fe(II) ions (25 μM, 100 μM, 400 μM).

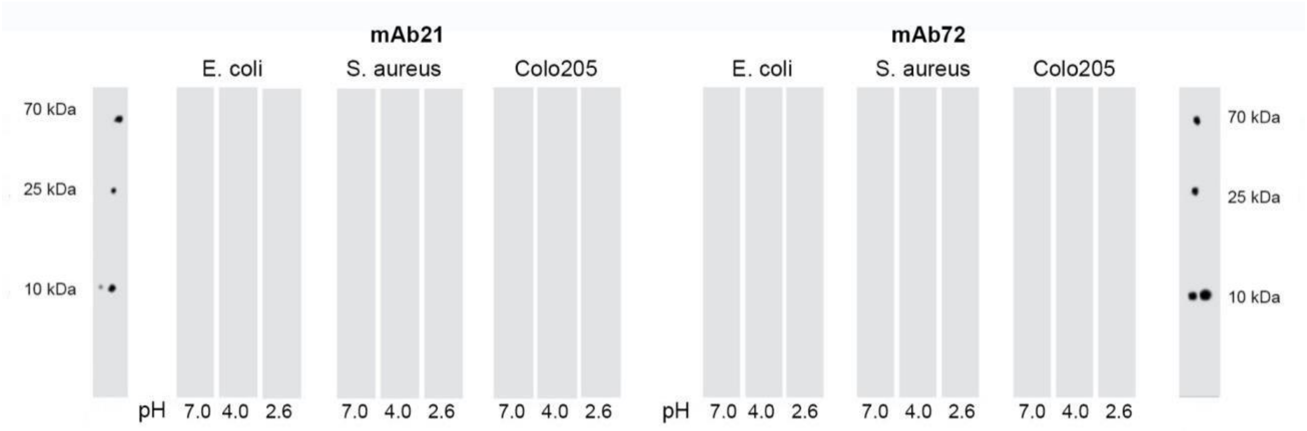


**Figure S3.** Western blot analysis of monoclonal IgA (mAb21 and mAb27) with extracts of *E. coli*, S. *aureus* и *Colo 205.* Antibodies was exposed to acid buffers (pH 4.0 and pH 2.6).

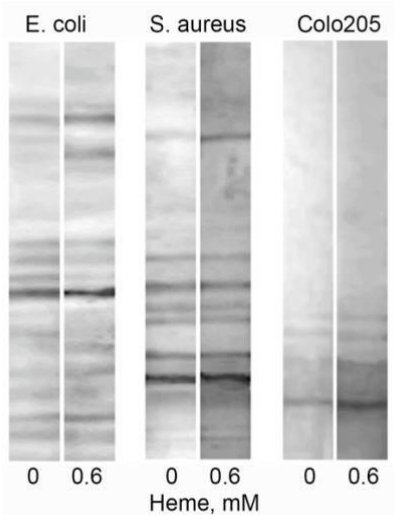


**Figure S4.** Western blot analysis of the binding of pooled human serum IgA to antigens in extracts from *E. coli, S. aureus* and Colo 205. The immunoglobulin was exposed to pH 2.6 buffer only (see the left blots in the three groups marked with “*0”)* or to pH 2.6 buffer plus 0,6 µM of heme (right blots, marked with “0.6”).
